# Evolution of Plasticity in Response to Ethanol between Recently Separated Populations of *D. melanogaster* with Different Ecological Histories

**DOI:** 10.1101/2023.06.22.546145

**Authors:** George Boateng-Sarfo, Franz Scherping, Murad Mammadov, Sarah Signor

## Abstract

The evolution of phenotype plasticity by genetic accommodation was theorized but has not been extensively demonstrated. Under this model of evolution when a population encounters a new environment there are widely variable responses amongst different genotypes, which are then pruned by selection into a single adaptive response. Because of the requirement to replicate genotypes, testing this predication requires inbred lines as well as populations that are both adapted and not adapted to a resource. We previously demonstrated that *D. melanogaster* adapted to ethanol through genetic accommodation using *D. simulans* as an ancestral proxy lineage. However, we wondered how generalizable these results were. Using a new population of *D. melanogaster* from France and an ancestral range population from Zambia we demonstrated here that the Zambian population is not adapted to ethanol and that the French population has evolved ethanol resistance through genetic accommodation. We also investigated alternative splicing in response to ethanol and find that gene expression and splicing appear to evolve independently of one another, and that the splicing response to ethanol is largely distinct between populations. Thus we have independently replicated evidence for evolution by genetic accommodation in *D. melanogaster*, suggesting that the evolution of plasticity may be an important contributor to the ability to exploit novel resources.

## Introduction

The adaptive evolution of phenotype plasticity is predicted to occur through the process of genetic accommodation. There are two predictions relating to the process of genetic accommodation in populations, based off of the stage of adaptation. First, a population must encounter a novel environment, and given that they are not adapted to that environment, there will be considerable variation between individuals in how they respond to the environment (Figure 1)^1–6^. At this stage the set of plastic responses in the population have not been selected on, thus they may be adaptive, deleterious, or neutral^7–10^. Adaptation to the novel environment will entail pruning of the plastic responses in the population to a single response that maximizes fitness^11^. However, it must also be adaptive to retain a plastic response rather than a fixed change^12–16^.

**Figure 1:**
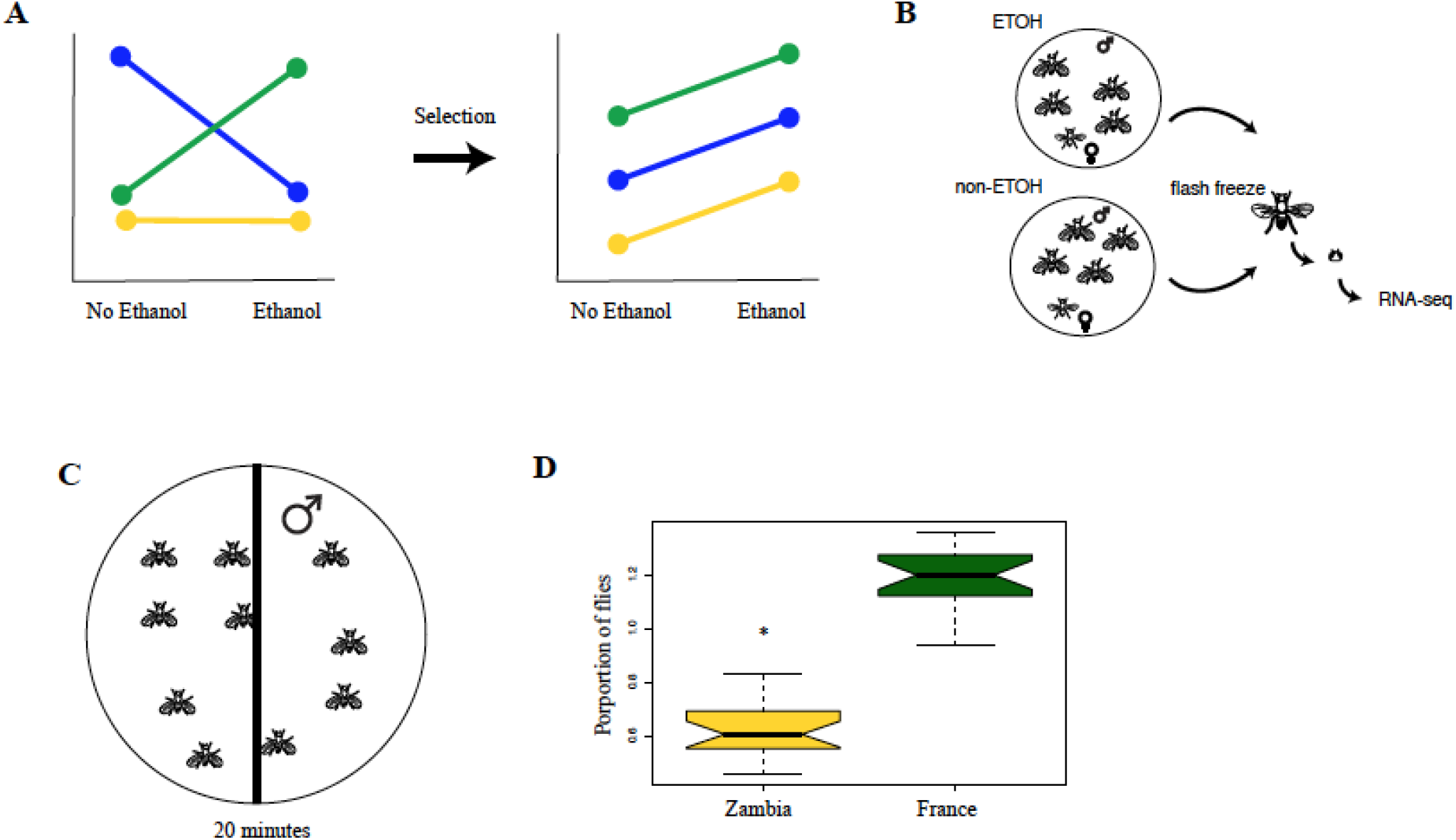
A, An illustration of the predictions for phenotypic plasticity and what was shown in *D. melanogaster* and *D. simulans*. This would be for one trait, for example a gene whose expression changes in response to ethanol. If the species is not adapted to ethanol (D. simulans) you would expect to see that different genotypes respond differently, that is, genotype by environment interactions. This is what we found in D. simulans. If the species is adapted to ethanol, you would expect each genotype to respond the same way (right). This is what we found in cosmopolitan D. melanogaster from California. What we are testing here is whether this prediction holds true between cosmopolitan D. melanogaster (France) and those from the native range of D. melanogaster (Zambia) who are not adapted to ethanol. B. An illustration of the environment that each Drosophila male was exposed to during the experiment. Each chamber contained ∼24 male flies and several females. The males were collected and flash frozen after 30 minutes. After flash freezing, their heads were isolated for RNA-seq. This was done for three genotypes each of Zambian D. melanogaster and French D. melanogaster. C. An example of the behavioral setup used to confirm that the two populations of D. melanogaster have different responses to ethanol. Every minute for twenty minutes the number of flies who crossed the midline of the plate were recorded. The substrate contains 15% ethanol. D. The proportion of flies that were not sedated and able to cross the midline of the plate was significantly higher in French populations of D. melanogaster.

**Figure 2:**
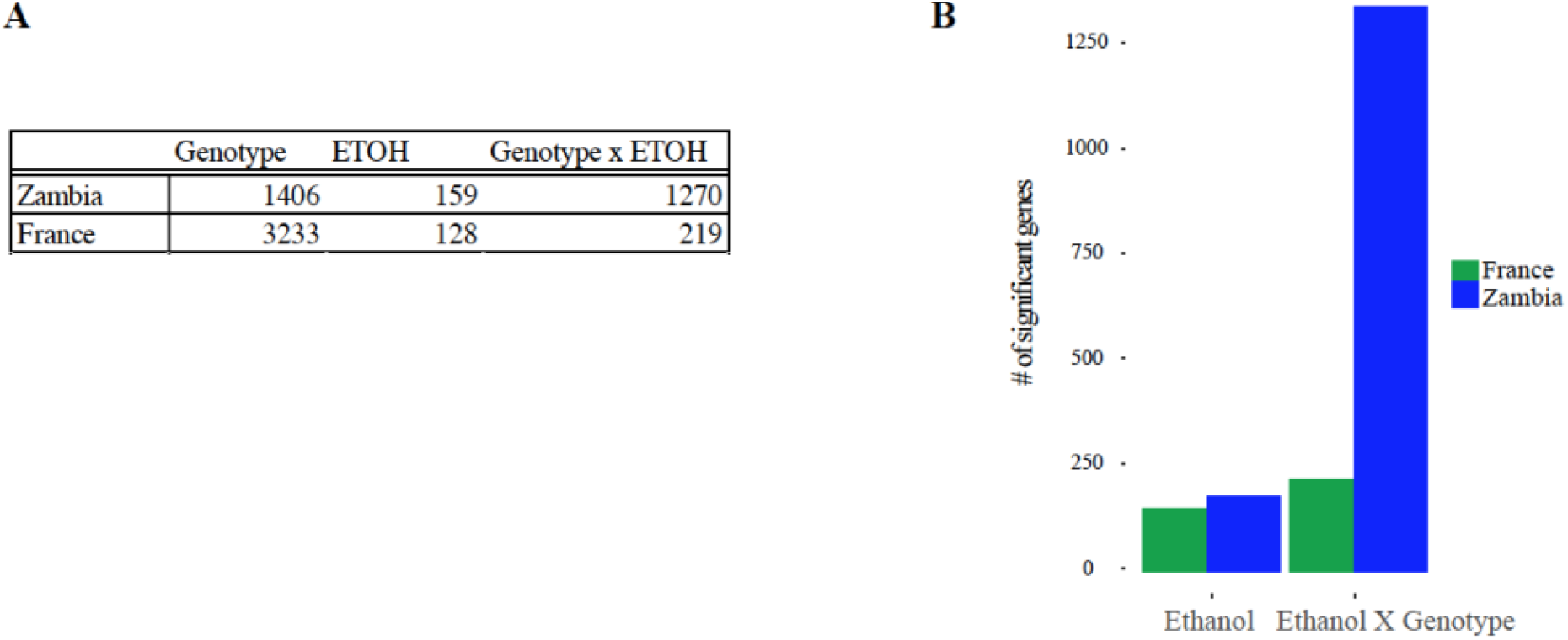
**A**. The number of genes that have a significant change in relative abundance for each component of variance in *D. melanogaster* from France and Zamiba. **B**. An illustration of the number of genes that change their expression in the presence of ethanol in each population of D. melanogaster. It is relatively similar for the effect of treatment, but in Zambian *D. melanogaster* the effect of genotype is very pronounced compared to cosmopolitan *D. melanogaster*.

Previously, Signor (2020) established that adaptation to ethanol in *D. simulans* and *D. melanogaster* met the expectations for genetic accommodation. Cosmopolitan populations of *D. melanogaster* are tolerant of ethanol and are found feeding and ovipositing on resources with ethanol concentrations greater than 8%^17,18^. Low tolerance to ethanol is the ‘ancestral state’, as most drosophilids are not tolerant to ethanol, and as such *D. simulans* avoids ethanol rich substrates^19–24^. Signor (2020) showed that four guidelines for establishing that the evolution of plasticity by genetic accommodation were met in this species pair^25,26^. First, the focal trait must be environmentally induced (ethanol) in a derived lineage (*D. melanogaster*) and an ancestral-proxy lineage (*D. simulans*). Second, cryptic genetic variation must be uncovered when the ancestral proxy lineage is exposed to the derived environment – we found that in *D. simulans* each genotype interacted very differently to ethanol compared to *D. melanogaster* were each genotype responded the same. This also relates to the third and fourth requirement, that the trait must show evidence of evolutionary change in the derived lineage and that the focal trait must exhibit evidence of adaptive refinement in the derived lineage. In summary, *D. simulans* exhibited extensive variation between genotypes in the plastic response to ethanol, evidence that cryptic genetic variation had not been removed by selection. Cosmopolitan *D. melanogaster* had a single plastic response to ethanol, evidence that selection has removed differences between genotypes.

Although this was strong evidence of evolution by genetic accommodation, and *D. simulans* does show the ancestral trait of low ethanol tolerance, we wondered how generalizable these results were and if we would find the same evidence if we compared cosmopolitan *D. melanogaster* to African *D. melanogaster*. African *D. melanogaster* in more remote locations (i.e. they do not show evidence of cosmopolitan admixture) still show the ancestral trait of low ethanol tolerance^27^. Demographic models suggest that *D. melanogaster* began its expansion out of Africa 10,000 years ago and colonized Europe approximately 2,000 years ago^28,29^. Thus this represents a recently evolved difference between these populations. If we found the same evidence - namely genotype specific plasticity in African *D. melanogaster* - between these more recently separated populations, this would be additional solid evidence for evolution by genetic accommodation.

We quantified gene and transcript level differential expression between ethanol exposed and non-ethanol exposed *D. melanogaster* from France and Zambia. We used three genotypes from each location to estimate the contribution of genotype to the response to ethanol treatment. What we found was that this system supports our conclusion of evolution by genetic accommodation. Both populations have approximately the same response to treatment overall, however in the Zambian population, each genotype responds differently to ethanol. In the French population, there is not a large contribution of genotype to the response to ethanol, suggesting that the plastic response has been pruned by selection. Furthermore, the genes that contribute to the response to treatment are consistent with our previous work on this system^6^. This continues to fit the model of evolution by genetic accommodation in this system, and is strong additional evidence for the importance of genetic accommodation in the evolution of plasticity.

## Methods

### Fly strains

Fly strains were generously provided by John Pool (UW-Madison) and originated either from France or Zambia^1,2^. The samples from Zambia were previously confirmed to be the most diverse among all the globally sampled strains, with minimal non-African admixture, suggesting they come from the ancestral range of *D. melanogaster*^1,2^.

### Fly collection

To minimize variation due to non-focal effects all collection populations were set up with 10 one day old individuals of each sex from a particular genotype. Flies were reared on a standard Bloomington medium at 25°C with a 12-h light/12-h dark cycle. After 8-9 days the vials were cleared and F1 males were collected for ethanol exposure assays and RNA-seq. 30 mated males were collected within three days of one another and subjected to assays within 3-5 days.

### Experimental Setup

The flies were sedated through exposure to cold for 10 mins and placed in exposure chambers with a paintbrush. Each chamber contained 5 ml of standard grapefruit fly media or media in which 15% of the water had been replaced by ethanol. They were allowed to acclimate for 10 min prior to timing the 30 minute exposure. This acclimation period is standard for behavioral assays, as it is long enough for the initial startle response to ethanol to have concluded, however here it was included to standardize data with past observations^3–8^. The assays were conducted within a 2-h window after dawn, the period in which the flies are most active^9–11^. Replicates were conducted randomly across days under standardized conditions (25°C, 70% humidity). At the conclusion of the assay the flies were flash frozen in liquid nitrogen. Frozen nested sieves were used to separate their bodies from their heads, limbs, and wings. Heads were collected for sequencing.

### Behavioral assays

Differences in the response to ethanol has been previously observed in African and cosmopolitan *D. melanogaster*^12–15^. However, we wanted to confirm that African populations had a lower tolerance for ethanol. A lower tolerance for ethanol would be indicated by a quicker progression through the euphoric stage of alcohol exposure to sedation^6,16^. The assays were conducted within a two-hour window after dawn, to standardize for the effect of circadian rhythms. Replicates were conducted randomly under standardized conditions (25 °C, 70% humidity). Approximately 30 male flies from a single genotype were sedated in a refrigerator for 10 minutes and then placed in a petri dish with 5 ml of grapefruit media in which 15% of the water had been replaced with ethanol. The petri dish was bisected by a black line, and every minute the flies were observed for 10 seconds. The number of flies that crossed the black line were recorded as a proxy for activity level, indicating that the flies were not sedated if they crossed the black line. This was done for ten minutes. Three replicates for each of the three genotypes was performed. Please note that a behavioral analysis of this exposure to ethanol has been published and shows evidence of intoxication as well as genotype-specific differences in the behavioral response to ethanol^6,30–32^.

### RNA sequencing

RNA was extracted from the heads of 30 male flies using the NucleoZol one phase RNA purification kit (Macherey-Nagel). Library preparation and sequencing was performed by BGI (Wuhan, China). The libraries were barcoded and pooled, and 2 million reads were generated per library on the illumina NextSeq. The data was demultiplexed prior to delivery.

### Differential Expression Analysis

Transcripts were quantified using the *D. melanogaster* reference transcriptome v.6.49 from Flybase and salmon v1.3.0^33^. The transcript quantification was imported into DESeq format with tximport and the *D. melanogaster* v6.49 GTF file was imported with the GenomicFeatures R package^34,35^. Differential expression was evaluated with DESeq2 and *p*-values were adjusted for multiple testing with Benjamini-Hochberg correction^36,37^.

### Alternative splicing analysis

Transcripts were mapped to the *D. melanogaster* v.6.49 genome with STAR aligner v.2.7.10a^38^. To quantify alternative splicing events we used rMATs (turbo) (v. 4.1.2)^39–41^ and the *D. melanogaster* annotation file v. 6.49 from Flybase^42^. We did not allow unannotated splice sites. We limited our analysis to an FDR <. 05 and ΔPSI > 0.1, including alternative 5’ splice sites, skipped exons, mutually exclusive exons, retained introns, and alternative 3’ splice sites. In general splice detection analysis does not include interaction terms, therefore this analysis was limited to the effect of ethanol in each population.

## Results

### Behavioral assays

*D. melanogaster* from Zambia had a significantly lower tolerance to ethanol than French *D. melanogaster* (Figure 1, two tailed t-test *p* <. 0001). Essentially the French *D. melanogaster* were able to stay in the euphoric stage of ethanol exposure without becoming sedated as the Zambian *D. melanogaster*. This confirms the expected difference in ethanol tolerance between African and cosmopolitan *D. melanogaster*. As such we know that cosmopolitan *D. melanogaster* are adapted to ethanol exposure and Zambian *D. melanogaster* are not.

### Changes in gene expression

Table 1 summarizes the number of genes which altered their expression in response to ethanol (Supplemental File 1 & 2). 159 genes change their expression in response to treatment in the Zambian *D. melanogaster*, while 128 change their expression in the French *D. melanogaster*. The most striking differences is in the interaction term between genotype and treatment, where cosmopolitan *D. melanogaster* exhibited only 219 changes while Zambian *D. melanogaster* had 1,270 genes which showed differential expression, a 6x increase. One explanation for the large effect of genotype in Zambian *D. melanogaster* compared to French *D. melanogaster* would be more variation attributable to genotype overall in Zambian *D. melanogaster*, particularly because the out of Africa expansion likely included a bottleneck^43–46^. However, this does not appear to be the case as in French *D. melanogaster* 3,233 genes are expressed differently between genotypes, and in Zambian *D. melanogaster* this number is 1,406 (Table 1). The disparity between the two populations of *D. melanogaster* is unique to categories that include an interaction with genotype.

### Core components of the response to ethanol

Genes that change their expression in both populations of *D. melanogaster* for any component of variance are likely to be core components of the response to ethanol. 149 genes had a response to ethanol in the Zambian and French populations of *D. melanogaster* (though not necessarily the same response), including *Drat, Pinocchio, cabut, sugarbabe*, and *Fatty Acid Synthase 2*. Compared to several other studies, four of the aforementioned genes are repeatedly implicated (*Drat, Pinocchio, cabut, sugarbabe*)^6,31,47–50^. In addition, in at least two other studies and in our dataset *FASN2, betaTub65B, AcCoAS, Pgd*, and *CG13607* were implicated in the response to ethanol^6,31,47–50^. This suggests that these are core components of the response to ethanol, given that in many studies overlap between gene expression datasets can be low. Interestingly the direction and magnitude of change of *Drat, Pinocchio, cabut*, and *sugarbabe* was the same between populations, suggesting they may be part of a conserved response to ethanol rather than part of the adaptive response in the French population.

### Alternative splicing

In general the splicing response to ethanol was larger in Zambian populations of *D. melanogaster*, with 88 SE (54 France), 57 A3S (32 France), 43 RI (25 France), and 41 A5S (31 France) (Table 2, Supplemental File 3 & 4). 34 genes and 32 events are shared between populations amongst all these categories, but of those only five are in the same direction, suggesting that the response to ethanol is quite distinct between populations (Table 2, Figure 3). This number is 56 genes if you consider across types of splicing. In the Zambian population some known candidate genes like *slowpoke, CASK*, and *dunce* are implicated in the splicing response, though only *slowpoke* was shown to be alternatively spliced^51–55^.

**Figure 3:**
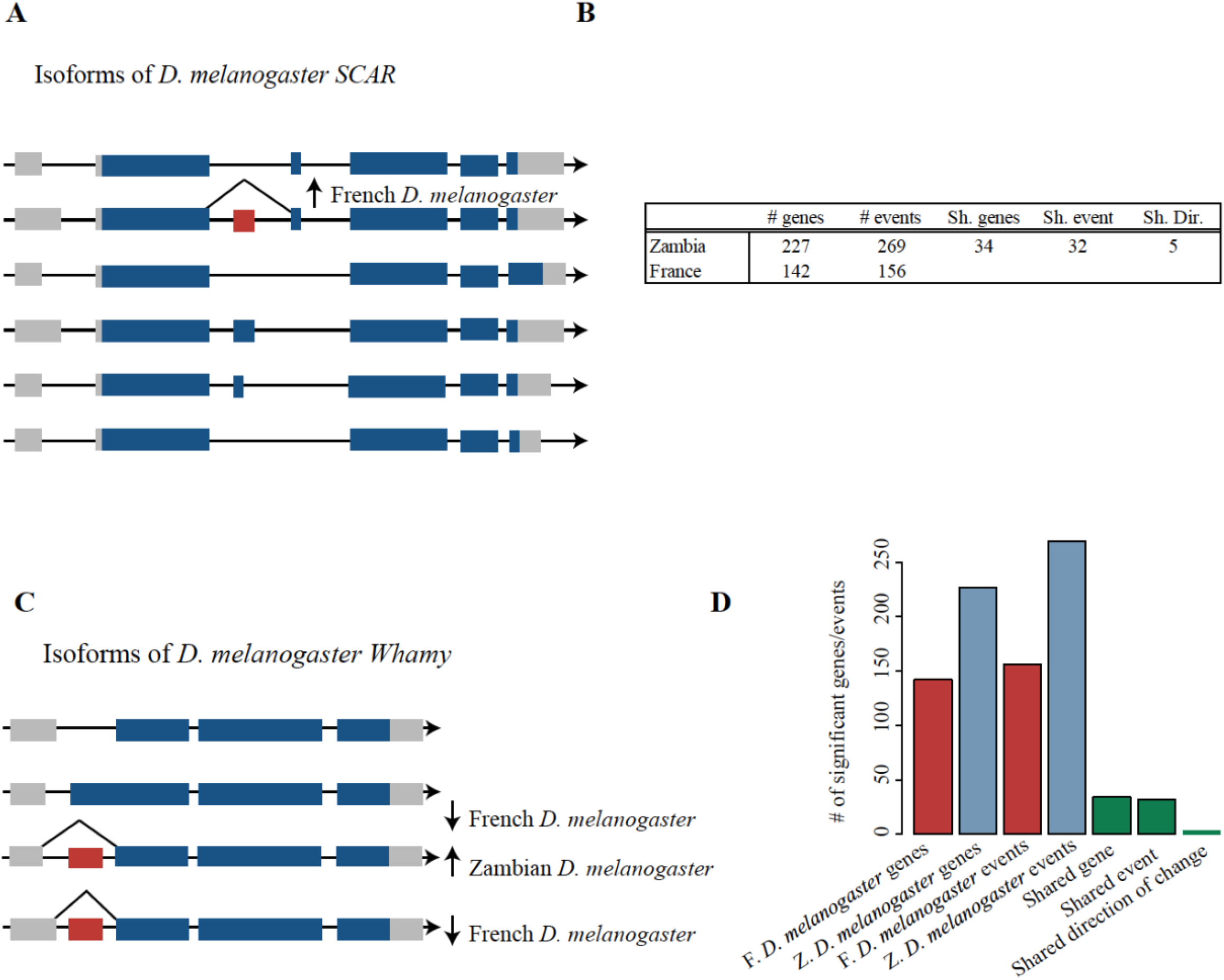
**A**. An example of the complex changes in isoform representation occurring in different populations of *D. melanogaster* for the gene *SCAR*. Grey boxes represent 3’ or 5’ UTRs, and the arrows on the left indicate the direction of transcription. Blue or red boxes indicate coding regions. This skipped exon increased in frequency in ethanol exposed French *D. melanogaster*. **B**. The total number of splicing differences for every category from each population. Sh. is the abbreviation for shared, dir. for direction. **C**. An example of changes in the representation of exons in the gene *Whamy*, which changed in different directions in each population. Inclusion of an alternative exon changed in two transcripts in French *D. melanogaster*, decreasing in frequency, while this exon increased in frequency in one transcript in Zambian *D. melanogaster*. **D**. This box plot illustrates the number splicing differences that was significant at p <. 05 and with a change in psi of at least 0.1 in each population of *D. melanogaster* (red and blue). Following that, shown in green, is the number of splicing events occurring in the same gene, the same event, and with the same direction of change.

In the adapted French population a number of genes known to interact with one another in muscle specification are implicated, including *spalt major, Myofilin, Stretchin-Mlck, frayed, myospheroid, SCAR, sallimus, tropomodulin, upheld*, and *wings upA* (Figure 3). Interestingly, *mef2* is thought to be a general regulator of ethanol sedation and is responsible for both muscle development and regulation of gene expression in neural tissue^56,57^. Indeed, *mef2* has a splicing difference in the Zambian population, and 39 genes in the Zambian population were also uncovered in a screen of *mef2* targets, including *Myosin heavy chain, shaking B*, and *dunce* ^57^. In the French population 33 genes were also implicated in this screen of *mef2* targets, including *fruitless, dachshund, Bruce, myospheroid*, and *frayed. Mef2* and *spalt major* are both myogenic regulators, but their relationship to each other is not clear. This raises the interesting possibility that the myogenic pathways may be an important part of the splicing response to ethanol. Overall overlap with previous work on alternative splicing in response to ethanol is low, with *Slowpoke binding protein, Syncrip, tweek, wings up A, midline fasciclin, Myosin heavy chain*, and *Rab3 interacting molecule* being found in at least one population here and previous work^6,31,58^.

### Core components of the splicing response

Despite the direction of change commonly being different, splicing responses in the same gene are likely part of the core response to ethanol. This includes known candidates such as *Syncrip* and *NFAT*, as well as new candidates such as *smooth, myospheroid*, and *suppressor of hairy wing. Myospheroid, NFAT, smooth, scalloped*, and *terribly reduced optic lobes* altered splicing in the same direction, suggesting they are not part of the adaptive response to ethanol. However, this is complicated, for example a skipped exon in *smooth* changed in opposite directions in the two populations, while 3’ alternative splicing changed in the same direction, suggesting there are likely still differences in isoform abundance at this gene. *Syncrip* was found in both populations changing in the opposite direction and was implicated in previous work in alternative splicing in response to ethanol^6,31^.

### Splicing and gene expression evolution are distinct

In the adapted French population no genes were shared between the gene expression and splicing response to ethanol. Between the splicing response to ethanol and the genotype by treatment gene expression differences five genes were shared. In the non-adapted Zambian population a single gene is shared between gene expression and splicing response to ethanol. Between the splicing response to ethanol and the genotype by treatment expression differences 19 genes were shared (out of more than a thousand that changed in the Zambian expression differences). Thus the splicing and gene expression differences in response to ethanol are largely distinct.

## Discussion

Previously it was shown that *D. simulans* and *D. melanogaster* met the predictions for evolution by genetic accommodation with regards to ethanol exposure, with *D. simulans* as the ancestral proxy population^6^. In *D. simulans* there was be abundant variation in how each genotype responds to ethanol, and in cosmopolitan *D. melanogaster* there was little variation (Figure 1A). However, it is possible that the observed pattern was isolated to this species pair and not a generalizable pattern. Here we have demonstrated that that is not in fact the case, and that genetic accommodation has occurred within the *D. melanogaster* lineage in the response to ethanol. Ancestral range *D. melanogaster* that are not adapted to ethanol show the same pattern of genotype-specific responses. Without selection on the response to ethanol, environmentally induced variants can accumulate as cryptic genetic variation and manifest as greater variation between genotypes^7,59–61^. Because ethanol is a patchy resource the response to ethanol is expected to be selected on as an optimal plastic response^12–16^.

Ancestral range and cosmopolitan *D. melanogaster* thus fit the criteria described previously for establishing evolution by genetic accommodation^25,26^. Using ancestral range *D. melanogaster* as an proxy lineage for ancestral *D. melanogaster* the focal trait can be environmentally induced by exposing them to ethanol. This exposure uncovers cryptic genetic variation – a large increase in genotype specific responses to ethanol. Further, while our previous work showed a loss of genotype specific response in Californian cosmopolitan *D. melanogaster*, this work confirmed this loss of genotype specific responses in a second French population of *D. melanogaster*^6^. Lastly, the focal trait shows evidence of adaptive refinement in cosmopolitan *D. melanogaster* – confirmed here with the increased resistance of French *D. melanogaster* and recorded elsewhere in terms of preferential oviposition in ethanol rich substrate^62,63^.

We also demonstrate here that gene expression and splicing have evolved independently, confirming that they have a separate genetic basis. This is consistent with recent predictions which suggest that separate responses for expression and splicing will be common and an important contributor to plasticity over short timescales^64^. The implication is that splicing and gene expression will also have distinct functions and affect different pathways, and indeed the two datasets contain genes with very different functions – for example the French splicing dataset contains many *mef2* interacting genes and genes thought to be involved in muscle/brain specification, while the gene expression datasets do not.

Previously we reported that in *D. simulans* the response to ethanol was enriched for non-protein coding genes as well as nested genes (genes in the introns of other genes). We hypothesized that cryptic genetic could variation affecting gene expression preferentially accumulate intronic non-protein coding genes due to lower selective constraint on expression. However, we did not find any enrichment in African *D. melanogaster* suggesting this is not a general feature of the accumulation of cryptic genetic variation. We also did not find a larger contribution of splicing in the adapted population, though we did find a larger contribution of splicing in California *D. melanogaster* compared to *D. simulans*. However, the method used to quantify splicing is very different between these manuscripts, as the previous paper used a more bespoke pipeline^6,31^. The splicing response was very distinct between populations, with only five genes changing in the same direction.

In our analysis of gene expression differences we uncovered some of the same genes which have been implicated in other studies including our own - *Drat, cabut, sugarbabe*, and *Pinocchio. cabut* and *Pinocchio* are also thought to be targets of *mef2*, potentially one of the core regulators of the response to ethanol^56,57^. While gene expression differences did not overlap considerably with the list of *mef2* targets, in the French population alternative splicing did, and contained many other myogenic genes that either interact with with *spalt major* or have been implicated in that pathway. While *mef2* and *spalt major* are referred to as myogenic genes *mef2* is required for mushroom body development, neuronal plasticity, and circadian rhythms^56,57,65–68^. *Spalt major* has a role in the brain as well, and is also a known target of *terribly reduced optic lobes* which also was implicated in splicing differences in the French population. Many of these genes appear to be transcription factors which may play a role in many essential functions, interacting with the same set of genes but deployed in different contexts. For example, *spalt major* is responsible for alternative splicing of *Myofilin* in developing muscles, and we see in this dataset changes in the splicing of both, but likely they are performing some kind of neural function^69^.

## Acknowledgements

Thanks to John Pool for generously providing the fly strains used in this study, and J. Saltz for feedback on the manuscript. Thanks to reviewer number two on our last reaction norm manuscript who convinced us to follow up on that study out of spite.

## Authors’ contributions

SS conceived the study, performed bioinformatics and drafted portions of the manuscript. GSB performed behavioral experiments and assisted in drafting the manuscript. FSB and MM performed experiments and collected material.

## Availability of data and materials

All data has been made available in the following repositories: The RNA-seq data has been made available under NCBI Bioproject number _.

## References

1 West-Eberhard MJ. Developmental plasticity and the origin of species differences. Proceedings of the National Academy of Sciences 2005;102 Suppl 1:6543–9. https://doi.org/10.1073/pnas.0501844102.

2 Ghalambor CK, Mckay JK, Carroll SP, Reznick DN. Adaptive versus non-adaptive phenotypic plasticity and the potential for contemporary adaptation in new environments. Functional Ecology 2007;21:394–407. https://doi.org/10.1111/j.1365-2435.2007.01283.x.

3 Robinson BW. Evolution of growth by genetic accommodation in Icelandic freshwater stickleback. Proceedings of the Royal Society B 2013;280:20132197–20132197. https://doi.org/10.1098/rspb.2013.2197.

4 Morris MRJ, Richard R, Leder EH, Barrett RDH, Aubin-Horth N, Rogers SM. Gene expression plasticity evolves in response to colonization of freshwater lakes in threespine stickleback. Molecular Ecology 2014;23:3226–40. https://doi.org/10.1111/mec.12820.

5 Schlichting CD, Wund MA. Phenotypic plasticity and epigenetic marking: an assessment of evidence for genetic accomodation. Evolution 2014;68:656–72. https://doi.org/10.1111/evo.12348.

6 Signor SA, Nuzhdin SV. Evolution of phenotypic plasticity in response to ethanol between sister species with different ecological histories (Drosophila melanogaster and D. simulans). BioRxiv 2019;3:72–27. https://doi.org/10.1101/386334.

7 Schlichting CD. Hidden reaction norms, cryptic genetic variation, and evolvability. Annals of the New York Academy of Sciences 2008;1133:187–203. https://doi.org/10.1196/annals.1438.010.

8 Gibson G. Decanalization and the origin of complex disease. Nature Reviews Genetics 2009;10:134–40. https://doi.org/10.1038/nrg2502.

9 Hayden EJ, Ferrada E, Wagner A. Cryptic genetic variation promotes rapid evolutionary adaptation in an RNA enzyme. Nature 2011;474:92–5. https://doi.org/10.1038/nature10083.

10 Paaby AB, Rockman MV. Cryptic genetic variation: evolution’s hidden substrate. Nature Reviews Genetics 2014;15:247–58. https://doi.org/10.1038/nrg3688.

11 Baldwin JM. A new factor in evolution. The American Naturalist 1896;30:441–51.

12 Via S, Lande R. Genotype-environment interaction and the evolution of phenotypic plasticity. Evolution 1985;39:505–22. https://doi.org/10.1111/j.1558-5646.1985.tb00391.x.

13 Guntrip J, Sibly RM. Phenotypic plasticity, genotype-by-environment interaction and the analysis of generalism and specialization in Callosobruchus maculatus. Heredity 1998;81:198– 204. https://doi.org/10.1046/j.1365-2540.1998.00354.x.

14 Lande R. Adaptation to an extraordinary environment by evolution of phenotypic plasticity and genetic assimilation. Journal of Evolutionary Biology 2009;22:1435–46. https://doi.org/10.1111/j.1420-9101.2009.01754.x.

15 Matzkin LM. Population transcriptomics of cactus host shifts in Drosophila mojavensis. Molecular Ecology 2012;21:2428–39. https://doi.org/10.1111/j.1365-294x.2012.05549.x.

16 Huang Y, Tran I, Agrawal AF. Does genetic variation maintained by environmental heterogeneity facilitate adaptation to novel selection? The American Naturalist 2016;188:27–37. https://doi.org/10.5061/dryad.kb830.

17 McKenzie JA, McKechnie SW. A comparative study of resource utilization in natural populations of Drosophila melanogaster and D. simulans. Oecologia 1979;40:. https://doi.org/10.1007/bf00345326.

18 Gibson J, May T, Wilks A. Genetic variation at the alcohol dehydrogenase locus in Drosophila melanogaster in relation to environmental variation: Ethanol levels in breeding sites and allozyme frequencies. Oecologia 1981;2:191–8. https://doi.org/10.1007/bf00540600.

19 Mercot H, Defaye D, Capy P, Pla E, David JR. Alcohol tolerance, ADH activity, and ecological niche of Drosophila species 1994;3:746–57. https://doi.org/10.2307/2410483.

20 Parsons PA, Stanley SM, Spence GE. Environmental ethanol at low concentrations: Longevity and development in the sibling species Drosophila melanogaster and D. simulans. Australian Journal of Zoology 1979;27:747–54. https://doi.org/10.1071/zo9790747.

21 Ziolo LK, Parsons PA. Ethanol tolerance, alcohol-dehydrogenase activity and Adh allozymes in Drosophila melanogaster. Genetica 1982;57:231–7. https://doi.org/10.1007/bf00056488.

22 Parsons PA. Ethanol utilization: threshold differences among six closely related species of Drosophila. Australian Journal of Zoology 1980;28:535–41. https://doi.org/10.1071/zo9800535.

23 Parsons PA, King SB. Ethanol: Larval discrimination between two Drosophila sibling species. Experientia 1977;33:898–9. https://doi.org/10.1007/bf01951269.

24 Parsons PA. Larval reaction to alcohol as an indicator of resource utilization differences between Drosophila melanogaster and D. simulans. Oecologia 1977;30:141–6. https://doi.org/10.1007/bf00345417.

25 Levis NA, Pfennig DW. Evaluating ‘Plasticity-first’ evolution in nature: key criteria and empirical approaches. Trends in Ecology & Evolution 2016;31:563–74. https://doi.org/10.1016/j.tree.2016.03.012.

26 Jones BM, Robinson GE. Genetic accommodation and the role of ancestral plasticity in the evolution of insect eusociality. Journal of Experimental Biology 2018;221:jeb153163–11. https://doi.org/10.1242/jeb.153163.

27 Sprengelmeyer QD, Pool JE. Ethanol resistance in Drosophila melanogaster has increased in parallel cold-adapted populations and shows a variable genetic architecture within and between populations. Ecol Evol 2021. https://doi.org/10.1002/ece3.8228.

28 Sprengelmeyer Q. The Population History of Drosophila melanogaster and the Evolution of Ethanol Tolerance and Body Size, Adaptive Traits. University of Wisconsin-Madison; 2021; 2021.

29 Sprengelmeyer QD, Mansourian S, Lange JD, Matute DR, Cooper BS, Jirle EV, et al. Recurrent collection of Drosophila melanogaster from Wild African environments and genomic insights into species history. Mol Biol Evol 2019;37:627–38. https://doi.org/10.1093/molbev/msz271.

30 Signor SA, Abbasi M, Marjoram P, Nuzhdin SV. Conservation of social effects (Ψ) between two species of Drosophila despite reversal of sexual dimorphism. Ecology and Evolution 2017;7:10031–41. https://doi.org/10.1002/ece3.3523.

31 Signor S, Nuzhdin S. Dynamic changes in gene expression and alternative splicing mediate the response to acute alcohol exposure in Drosophila melanogaster. Heredity 2018.

32 Signor SA, Abbasi M, Marjoram P, Nuzhdin SV. Social effects for locomotion vary between environments in Drosophila melanogaster females. Evolution 2017;71:1765–75. https://doi.org/10.1111/evo.13266.

33 Patro R, Duggal G, Love MI, Irizarry RA, Kingsford C. Salmon provides fast and bias-aware quantification of transcript expression 2017;14:417–9. https://doi.org/10.1038/nmeth.4197.

34 Soneson C, Love MI, Robinson MD. Differential analyses for RNA-seq: transcript-level estimates improve gene-level inferences. F1000research 2016;4:1521. https://doi.org/10.12688/f1000research.7563.2.

35 Lawrence M, Huber W, Pagès H, Aboyoun P, Carlson M, Gentleman R, et al. Software for computing and annotating genomic ranges. Plos Comput Biol 2013;9:e1003118. https://doi.org/10.1371/journal.pcbi.1003118.

36 Benjamini Y, Hochberg Y. Controlling the false discovery rate: a practical and powerful approach to multiple testing. Journal of the Royal Statistical Society, Series B n.d.;57:289–300. https://doi.org/10.2307/2346101.

37 Love MI, Huber W, Anders S. Moderated estimation of fold change and dispersion for RNA-seq data with DESeq2. Genome Biology 2014;15:31–21. https://doi.org/10.1186/s13059-014-0550-8.

38 Dobin A, Davis CA, Schlesinger F, Drenkow J, Zaleski C, Jha S, et al. STAR: ultrafast universal RNA-seq aligner. Bioinformatics 2013;29:15–21. https://doi.org/10.1093/bioinformatics/bts635.

39 Shen S, Park JW, Lu Z, Lin L, Henry MD, Wu YN, et al. rMATS: Robust and flexible detection of differential alternative splicing from replicate RNA-Seq data. Proc National Acad Sci 2014;111:E5593–601. https://doi.org/10.1073/pnas.1419161111.

40 Park JW, Tokheim C, Shen S, Xing Y. Identifying differential alternative splicing events from RNA sequencing data using RNASeq-MATS. Totowa, NJ: Humana Press; 2013. p. 171–9.

41 Shen S, Park JW, Huang J, Dittmar KA, Lu Z, Zhou Q, et al. MATS: a Bayesian framework for flexible detection of differential alternative splicing from RNA-Seq data. Nucleic Acids Res 2012;40:e61–e61. https://doi.org/10.1093/nar/gkr1291.

42 Gramates LS, Agapite J, Attrill H, Calvi BR, Crosby MA, Santos G dos, et al. FlyBase: a guided tour of highlighted features. Genetics 2022;220:iyac035. https://doi.org/10.1093/genetics/iyac035.

43 Li H, Stephan W. Inferring the demographic history and rate of adaptive substitution in Drosophila. PLoS Genetics 2006;2:e166. https://doi.org/10.1371/journal.pgen.0020166.

44 Ometto L, Glinka S, Lorenzo DD, Stephan W. Inferring the effects of demography and selection on Drosophila melanogaster populations from a chromosome-wide scan of DNA variation. Molecular Biology and Evolution 2005;22:2119–30. https://doi.org/10.1093/molbev/msi207.

45 Haddrill PR, Thornton KR, Charlesworth B, Andolfatto P. Multilocus patterns of nucleotide variability and the demographic and selection history of Drosophila melanogaster populations. Genome Research 2005;15:790–9. https://doi.org/10.1101/gr.3541005.

46 Li YJ, Satta Y, Takahata N. Paleo-demography of the Drosophila melanogaster subgroup: application of the maximum likelihood method. Genes & Genetic Systems 1999;74:117–27.

47 Morozova TV, Mackay TFC, Anholt RRH. Transcriptional networks for alcohol sensitivity in Drosophila melanogaster 2011;187:1193–205. https://doi.org/10.1534/genetics.110.125229.

48 Morozova T, Anholt R, Mackay TF. Transcriptional response to alcohol exposure in Drosophila melanogaster. Genome Biology 2006;7:R95. https://doi.org/10.1186/gb-2006-7-10-r95).

49 Kong EC, Allouche L, Chapot PA, Vranizan K, Moore MS, Heberlein U, et al. Ethanol-regulated genes that contribute to ethanol sensitivity and rapid tolerance in Drosophila. Alcoholism: Clinical and Experimental Research 2010;34:302–16. https://doi.org/10.1111/j.1530-0277.2009.01093.x.

50 Ghezzi A, Krishnan HR, Lew L, Prado FJ, Ong DS, Atkinson NS. Alcohol-induced histone acetylation reveals a gene network involved in alcohol tolerance. PLoS Genetics 2013;9:e1003986–14. https://doi.org/10.1371/journal.pgen.1003986.

51 Cowmeadow RB, Krishnan HR, Ghezzi A, Al’Hasan YM, Wang YZ, Atkinson NS. Ethanol Tolerance caused by slowpoke induction in Drosophila. Alcoholism: Clinical and Experimental Research 2006;30:745–53. https://doi.org/10.1111/j.1530-0277.2006.00087.x.

52 Cowmeadow RB, Krishnan HR, Atkinson NS. The slowpoke gene is necessary for rapid ethanol tolerance in Drosophila. Alcoholism: Clinical and Experimental Research 2005;29:1777–86. https://doi.org/10.1097/01.alc.0000183232.56788.62.

53 Rodan AR, Rothenfluh A. The Genetics of Behavioral Alcohol Responses in Drosophila. vol. 91. Elsevier Inc.; 2010.

54 Xu S, Chan T, Shah V, Zhang S, Pletcher SD, Roman G. The propensity for consuming ethanol in Drosophila requires rutabaga adenylyl cyclase expression within mushroom body neurons. Genes, Brain, and Behavior 2012;11:727–39. https://doi.org/10.1111/j.1601-183x.2012.00810.x.

55 Park A, Ghezzi A, Wijesekera TP, Atkinson NS. Genetics and genomics of alcohol responses in Drosophila. Neuropharmacology 2017:1–52. https://doi.org/10.1016/j.neuropharm.2017.01.032.

56 Schmitt RE, Shell BC, Lee KM, Shelton KL, Mathies LD, Edwards AC, et al. Convergent evidence from humans and Drosophila melanogaster implicates the transcription factor MEF2B/Mef2 in alcohol sensitivity. Alcohol Clin Exp Res 2019;43:1872–86. https://doi.org/10.1111/acer.14138.

57 Talikoti A. Identifying Genes Downstream of Mef2 that Influence Ethanol Sedat.pdf. Virginia Commonwealth University; 2021; 2021.

58 Petruccelli E, Feyder M, Ledru N, Jaques Y, Anderson E, Kaun KR. Alcohol activates Scabrous-Notch to influence associated memories. Neuron 2018;100:1209-1223.e4. https://doi.org/10.1016/j.neuron.2018.10.005.

59 Rutherford SL. From genotype to phenotype: buffering mechanisms and the storage of genetic information. BioEssays 2000;22:1095–105. https://doi.org/10.1002/1521-1878(200012)22:12<

60 Gibson G, Dworkin I. Uncovering cryptic genetic variation. Nature Reviews Genetics 2004;5:681–90. https://doi.org/10.1038/nrg1426.

61 Hermisson J, Wagner GP. The population genetic theory of hidden variation and genetic robustness. Genetics 2004;168:2271–84. https://doi.org/10.1534/genetics.104.029173.

62 Milan NF, Kacsoh BZ, Schlenke TA. Alcohol consumption as self-medication against blood-borne parasites in the fruit fly. Current Biology 2012;22:488–93. https://doi.org/10.1016/j.cub.2012.01.045.

63 Pohl JB, Baldwin BA, Dinh BL, Rahman P, Smerek D, Prado FJ, et al. Ethanol preference in Drosophila melanogaster is driven by its caloric value. Alcoholism: Clinical and Experimental Research 2012;36:1903–12. https://doi.org/10.1111/j.1530-0277.2012.01817.x.

64 Verta J-P, Jacobs A. The role of alternative splicing in adaptation and evolution. Trends Ecol Evol 2022;37:299–308. https://doi.org/10.1016/j.tree.2021.11.010.

65 Crittenden JR, Skoulakis EMC, Goldstein ES, Davis RL. Drosophila mef2 is essential for normal mushroom body and wing development. Biol Open 2018;7:bio035618. https://doi.org/10.1242/bio.035618.

66 Adhikari P, Orozco D, Randhawa H, Wolf FW. Mef2 induction of the immediate early gene Hr38/Nr4a is terminated by Sirt1 to promote ethanol tolerance. Genes, Brain, and Behavior 2018:e12486. https://doi.org/10.1111/gbb.12486.

67 Lin X, Shah S, Bulleit RF. The expression of MEF2 genes is implicated in CNS neuronal differentiation. Mol Brain Res 1996;42:307–16. https://doi.org/10.1016/s0169-328x(96)00135-0.

68 Sivachenko A, Li Y, Abruzzi KC, Rosbash M. The transcription factor Mef2 links the Drosophila core clock to Fas2, neuronal morphology, and circadian behavior. Neuron 2013;79:281–92. https://doi.org/10.1016/j.neuron.2013.05.015.

69 Spletter ML, Barz C, Yeroslaviz A, Schönbauer C, Ferreira IRS, Sarov M, et al. The RNA-binding protein Arrest (Bruno) regulates alternative splicing to enable myofibril maturation in Drosophila flight muscle. Embo Rep 2015;16:178–91. https://doi.org/10.15252/embr.201439791.

